# Genome-scale fitness profile of Caulobacter crescentus grown in natural freshwater

**DOI:** 10.1101/279711

**Authors:** Kristy L. Hentchel, Leila M. Reyes Ruiz, Patrick D. Curtis, Aretha Fiebig, Maureen L. Coleman, Sean Crosson

**Author notes:** To whom correspondence should be addressed; 929 E. 57th St. Chicago, IL 60637, USA.

## Abstract

Bacterial genomes evolve in complex ecosystems and are best understood in this natural context, but replicating such conditions in the lab is challenging. We used transposon sequencing to define the fitness consequences of gene disruption in the bacterium *Caulobacter crescentus* grown in natural freshwater, compared to axenic growth in common laboratory media. Gene disruptions in amino acid and nucleotide biosynthesis pathways and in metabolic substrate transport machinery impaired fitness in both lake water and defined minimal medium relative to complex peptone broth. Fitness in lake water was enhanced by insertions in genes required for flagellum biosynthesis and reduced by insertions in genes involved in biosynthesis of the holdfast surface adhesin. We further uncovered numerous hypothetical and uncharacterized genes for which disruption impaired fitness in lake water, defined minimal medium, or both. At the genome scale, the fitness profile of mutants cultivated in lake water was more similar to that in complex peptone broth than in defined minimal medium. Microfiltration of lake water did not significantly affect the terminal cell density or the fitness profile of the transposon mutant pool, suggesting that *Caulobacter* does not strongly interact with other microbes in this ecosystem on the measured timescale. Fitness of select mutants with defects in cell surface biosynthesis and environmental sensing were significantly more variable in lake water than in defined medium, presumably owing to day-to-day heterogeneity in the lake environment. This study reveals genetic interactions between *Caulobacter* and a natural freshwater environment, and provides a new avenue to study gene function in complex ecosystems.

## INTRODUCTION

Environments inhabited by microbial cells have significant microscale heterogeneity. This is well recognized in biofilms, soils, and host-associated habitats (1-3). Free-living aquatic bacteria often encounter chemical gradients that can appear as ephemeral patches, arising from algal exudates, sinking particles, or lysis events (4, 5) and they may have to cope with prolonged periods of nutrient scarcity. In addition, these bacteria face interspecies interactions, protistan predators, and viruses, as well as fluctuations in physical conditions such as temperature and light. These biotic and abiotic factors have driven myriad adaptations that enable survival and reproduction in natural environments.

In contrast with this natural complexity, studies on microbial physiology and gene function have traditionally relied on simplified experimental conditions. Thus it is not surprising that a large fraction of bacterial genes remain uncharacterized. Recently developed transposon sequencing (Tn-Seq) approaches (6) now enable rapid phenotypic assessment of thousands to millions of distinct mutant strains, and these methods have been used to interrogate gene function in a variety of in vitro and host-associated conditions (7, 8). More recently, transposon mutagenesis approaches in which each transposon carries a unique 20-bp barcode sequence have been developed (9); each insertion site is associated with a short barcode, and the abundance of all mutant strains in the pool can be assessed by simple amplicon sequencing.

Here, we used a barcoded Tn-Seq approach to identify genes affecting fitness in *Caulobacter crescentus* strain CB15, cultivated in natural freshwater from Lake Michigan, Illinois, USA. As a well-characterized and genetically tractable bacterium originally isolated from a pond in California in 1960 (10), this strain is well suited for this study. Briefly, *C. crescentus* is among a group of dimorphic prosthecate (i.e. stalked) bacteria that attach to surfaces, often forming epibiotic interactions with algae and plant material (11). More broadly, members of the genus *Caulobacter* are common in soil ecosystems, where they likely play an important role in plant matter decomposition (12). In aquatic systems, *Caulobacter* interactions with substrates contribute to biopolymer mineralization, and have been proposed to enhance productivity of aquatic ecosystems (11, 13). However, *C. crescentus* (hereafter referred to simply as *Caulobacter*) is typically grown in an artificial medium consisting of dilute peptone and yeast extract (PYE) or in a defined medium consisting of mineral salts and a single carbon source such as xylose (M2X) (14), neither of which adequately represents natural freshwater. PYE is replete with peptides, amino acids, and a range of carbon sources, while M2X requires *Caulobacter* to synthesize all cellular building blocks from salts and a simple sugar. Natural freshwaters, by contrast, contain an undefined, complex mixture of organic and inorganic nutrient sources (15). In many freshwater systems, essential nutrients including phosphorus and labile carbon do not accumulate to high concentrations (16, 17). We predicted that genes that are dispensable in PYE (18) or M2X medium would be important for fitness in complex natural freshwater, and that these genes would offer insights into *Caulobacter* physiology in a *bona fide* freshwater system.

## MATERIALS AND METHODS

### Bacterial strains and primers

Strains and primers used in this study are listed in **Table S1**. All primers were synthesized by Integrated DNA Technologies (Coralville, IA, USA).

### Growth media

*. Caulobacter crescentus* strain CB15 (10) was grown in PYE medium [0.2% peptone (Fisher Scientific), 0.1% yeast extract (Fisher Scientific), 0.5 mM MgSO4, 0.5 mM CaCl2] or M2X minimal defined medium [6.1 mM Na2HPO4, 3.9 mM KH2PO4, 9.3 mM NH4Cl, 0.5 mM MgSO4, 10 μM FeSO4 (EDTA chelate; Sigma Chemical Co.), 0.25 mM CaCl2] supplemented with 0.15% xylose (14). *Escherichia coli* strains were grown in LB broth (1% peptone, 0.5% yeast extract, 0.5% NaCl). Solid growth media included 1.5% agar.

### Lake water collection

Water from Lake Michigan was collected at Promontory Point, Chicago, Illinois, USA (Latitude: 41.794, Longitude: -87.579), on four dates in 2016 (Nov 30, Dec 6, Dec 9, and Dec 12). We measured water temperature, phosphate and nitrate/nitrite level (Aquacheck Water Quality Test Strips; Hach), and pH (pH indicator strips, Fisher Scientific) at the time of collection (**Table 1)**. Lake water was filtered using Nalgene™ Rapid-Flow™ Sterile Disposable 0.1 μm Filter Units with PES Membrane (Thermo Scientific).

**Table 1:** Analysis of Lake Michigan water used for barcoded Tn-Himar fitness experiments

|  |  |  |  |  |
| --- | --- | --- | --- | --- |
| Water collection<sup>a</sup> | Nov. 30 | Dec. 6 | Dec. 9 | Dec. 12 |
| Date of experiment | Dec. 2 | Dec. 6 | Dec. 9 | Dec. 12 |
| Water temperature (°C) | 7.5 | 7 | 3 | 2 |
| Air temperature (°C) | 7.8 | 4.4 | -3.3 | -7.2 |
| pH | 5.8 | 5.8 | 5.8 | 5.8 |
| Phosphate | 5 ppm | 5 ppm | 5 ppm | 5 ppm |
| Nitrate/Nitrite<sup>b</sup> | undetectable | undetectable | undetectable | undetectable |

<sup>a</sup>Water collection occurred in 2016 at Promontory Point, Hyde Park, Chicago, Illinois, USA.
<sup>b</sup>Limit of detection: Nitrite 0.15 mg/L, Nitrate 1 mg/L

### Measurement of Caulobacter growth in lake water

Colonies of *Caulobacter* were inoculated into 2 mL of PYE in glass culture tubes (13 × 100 mm) and grown overnight at 30°C with shaking at 200 rpm, for a total of five biological replicates. At saturation, 1 mL of culture was centrifuged at 8,000 ×*g* and washed twice in 1 mL of filtered lake water (see **Figure 2** for information on lake water). The washed pellet was resuspended in filtered lake water to a final OD660 of 0.1, and 0.5 μL (approximately 0.5–1 × 105 cells) was inoculated into 5 mL of filtered lake water in a glass culture tube (20 mm × 150 mm) in technical duplicate. Cultures were grown at 30°C with shaking at 200 rpm. To monitor growth, 20 μL of culture was removed at various time points, serially diluted, and titered onto PYE agar plates, which were incubated at 30°C for 2 days. Growth was monitored by enumeration of colony forming units (CFUs).

### Construction of barcoded Tn-Himar mutant library

The recipient strain (*Caulobacter*) was grown overnight in 2 mL of PYE at 30°C with shaking at 200 rpm. This starter culture was used to inoculate 20 mL of PYE and grown at 30°C overnight with shaking at 200 rpm until saturated. The donor *E. coli* strain (APA752, gift from Adam Deutschbauer, University of California-Berkeley, USA), carrying the pKMW3 (kanamycin resistant) Himar transposon vector library (9), was inoculated into 20 mL of LB containing kanamycin (30 μg mL^-1^) and diaminopimelate (DAP; 300 μM) and grown overnight at 37°C with shaking at 200 rpm; the *E. coli* Himar donor strain is a DAP auxotroph, and thus requires addition to the medium. To conjugate the barcoded transposon pool into *Caulobacter*, the recipient strain and donor strains were each centrifuged at 8,000 ×*g* for 2 min and resuspended in a total volume of 500 μL of PYE medium. The cultures were combined at a 10:1 ratio of recipient to donor and mixed by gentle pipetting. The mixed culture was centrifuged again at 8,000 ×*g*, and the supernatant decanted. The cells were resuspended in 30 μL of PYE, spotted onto a PYE agar plate containing diaminopimelate (300 μM), and incubated overnight at 30°C. After growth, the mating spot was scraped from the plate and resuspended in 6.5 mL of PYE. This suspension was spread evenly (500 μL per plate) over 14 large (150 × 15 mm) PYE agar plates containing 25 μg mL^-1^ kanamycin and incubated for approximately 3 days at 30°C. Cells were harvested from all the plates and inoculated into 400 mL of PYE containing 5 μg mL^-1^ kanamycin. This cell mixture was grown at 30°C with shaking at 200 rpm for three doublings. Cells were centrifuged at 8,000 ×*g*, resuspended in 70 mL of PYE containing 15% glycerol, and stored as 1 mL aliquots at -80°C.

### Mapping of the sites of Tn-Himar insertion in the Caulobacter BarSeq library

(see Fig. S1 for graphical overview). Genomic DNA was extracted using guanidium thiocyanate as previously described (19). The DNA was sheared (~300 bp fragments), cleaned with a standard bead protocol, end-repaired and A-tailed, and a custom double-stranded Y adapter was ligated. The custom adapter was prepared by annealing Mod2_TS_Univ and Mod2_TruSeq (**Table S1**) as described (9). The sheared fragments containing transposons were enriched by PCR using the primers Nspacer_BarSeq_pHIMAR and P7_MOD_TS_index1 using GoTaq^®^ Green Master Mix according to the manufacturer’s protocol in a 100-μL volume with the following cycling conditions: 94°C for 2 min, 25 cycles at 94°C for 30 s, 65°C for 20 s, and 72°C for 30 s, followed by a final extension at 72°C for 10 min. After a second bead cleanup, the *Caulobacter* library was sequenced using a standard Illumina sequencing primer on an Illumina HiSeq2500 at the University of Chicago Genomics Facility with a 150-bp single-end read. The locations of Himar transposon insertions were aligned and mapped using BLAT (20), and unique barcode sequences were associated with their corresponding genome insertion location using a custom Perl script (MapTnSeq.pl). Sets of barcodes that consistently map to one location in the genome were identified using a custom Perl script (DesignRandomPool.pl). This ensures that each unique barcode is properly assigned to a single insertion site. These scripts have been described by Wetmore and colleagues (9) and are available at https://bitbucket.org/berkeleylab/feba. For all analyses, reads were mapped to the *C.crescentus* NA1000 genome (accession CP001340) (21), which is more comprehensively annotated (22) than the highly-related CB15 parent strain.

### Cultivation of the Tn-Himar library

An aliquot of the *Caulobacter* library (2 mL) from a glycerol stock was inoculated into 18 mL of PYE, split into two tubes (20 × 150 mm) with 10 mL each, and grown in a cell culture roller drum (Fisher Scientific) at 30°C for 4 h. The tubes were then moved to a 30°C incubator with shaking at 200 rpm for an additional 2 h. Cultures were combined and centrifuged for 20 min at 3,000 ×*g* at 4°C. The cell pellet was resuspended and washed in 10 mL of filtered lake water, and centrifuged again at 3,000 ×*g* for 20 min at 4°C. The resulting pellet was resuspended in 5 mL of filtered lake water, and the OD660 measured. Flasks containing filtered or unfiltered lake water (7.5 L total volume per condition, divided over 3 flasks) were inoculated with the washed library with the aim of an initial starting concentration of approximately 2.5 × 10^7^ total cells per flask (**Fig. S2**). Flasks were incubated at 30°C with shaking at 150 rpm. At 0 and 64 h, an aliquot of culture was removed from each flask for CFU enumeration on PYE agar plates (**Fig. S3**). After ~64 h of growth, cells from all three flasks were collected by filtration using an Express Plus Membrane 0.22 μm filter (Millipore). Filters were stored at -80°C until needed. To mimic saturating conditions with the same number of doublings in defined M2X and complex PYE laboratory medium as in lake water, we inoculated cultures at a concentration that after five doublings (the estimated number of doublings in lake water), the cultures reached saturation. Cells were pelleted at 10,000 ×g for 1 min and stored at -20°C. Genomic DNA from all samples was extracted using guanidium thiocyanate as previously described (19), with the exception that the lake water samples were lysed directly from the filters they were collected on. DNA quality and quantity was measured using a NanoDrop^OneC^ (Thermo Scientific).

### Amplification and sequencing of Tn-Himar barcodes

PCR amplification for each sample was performed as previously described (9) (**Fig. S1**) using a standard reaction protocol for Q5 DNA polymerase (New England BioLabs) with the primers BarSeq_P1 and 1 of 16 forward primers (BarSeq_P2_IT001 to BarSeq_P2_IT016; **Table S1**) containing unique 6-bp TruSeq indexes that were sequenced using a separate index primer. Cycling conditions were as follows: 98°C for 4 min followed by 25 cycles of 30s at 98°C, 30s at 55°C, and 30s at 72°C, followed by a final extension at 72°C for 5 min. PCR products were purified using GeneJET PCR Purification Kit (Thermo Scientific). Purified samples were run on a 2.5% agarose gel to confirm correct product size (~200 bp). A total of 10 μL per purified PCR product was pooled, assessed for quality, and quantified using a Bioanalyzer. The amplified barcodes from the reference (PYE) and treatment (M2X, unfiltered lake water, and filtered lake water) were sequenced on an Illumina HiSeq4000 at the University of Chicago Genomics Facility, multiplexing all 16 samples in one lane with 50-bp single-end reads. All sequence data have been deposited in the NCBI Sequence Read Archive under BioProject accession PRJNA429486; BioSample accession SAMN08348121; SRA accession SRP128742.

### Analysis of Tn-Himar strain fitness

We followed the fitness calculation protocol of Wetmore and colleagues (9), using scripts available at https://bitbucket.org/berkeleylab/feba. Briefly, the total count of each barcode in each sample was calculated using a Perl script (MultiCodes.pl) and, from this table of barcodes, strain fitness was calculated using an R script (FEBA.R). The fitness of each strain was calculated as a normalized log2 ratio of barcode counts in the treatment sample to counts in the PYE reference sample. The fitness of genes was calculated as the weighted average of strain fitness values, the weight being inversely proportional to a variance metric based on the total number of reads for each strain; this weighting is fully described by Wetmore and colleagues (9). Successful gene fitness calculations required at least 3 reads per strain and 30 reads for each of the 16 samples. Insertions in the first 10% or last 10% of a gene were not considered in gene fitness calculations. The complete data set of fitness values for each condition is listed in **Table S2**.

To assess the distribution of fitness scores, we calculated the standard deviation for each condition using the frequency distribution of the mean fitness value of each gene (filtered lake water = 0.41, unfiltered lake water = 0.40, defined medium = 1.1). When the outlier region of the defined medium dataset (< -2.5) was removed, the calculated standard deviation was 0.36; therefore, a standard deviation of 0.4 was chosen and applied to all conditions. Genes with a mean fitness value approximated at ± 3σ from the mean (less than -1.2 and greater than +1.2) were selected for further examination. We also examined t-values, the fitness value of a gene divided by a variance metric, based on the total number of reads for each gene (as previously described (9)), to provide a metric to assess the significance of fitness values (**Table S3**).

To identify genes showing differential fitness across lake water samples, we fit a linear model with two factors, sampling day and filtration treatment (filtered or unfiltered). The model was implemented using the functions *lmfit, eBayes, and topTable* in the R package *limma* (23). Genes were identified as having differential fitness across either sampling days or filtration treatment, with a false discovery rate threshold of 0.05.

### Analysis of Caulobacter Tn5-seq fitness. A Caulobacter

Tn5 insertion library containing an estimated 3 × 10^5^ clones was constructed as previously described (24). The lake water fitness experiment for the Tn5 library (from Lake Michigan water collected in April 2016) was performed similarly to the Tn-Himar library experiments with the following modifications: A total of 200 μL of the *Caulobacter* Tn5 library was inoculated into 20 mL of PYE for the initial outgrowth for 5 h, which was then inoculated into 2 L for the PYE and unfiltered lake water treatments, and 2 replicates of 2 L each for filtered lake water. Lake water cultures were harvested by filtration after 60 h of growth, and the PYE condition was filtered after 12 h to approximate the same number of doublings. However, our PYE cultures achieved over 6 doublings, versus 4 doublings for lake water.

A nested PCR approach was used to specifically amplify transposon-containing DNA fragments for sequencing. A low cycle PCR amplification for each sample was first performed using a standard reaction protocol for KOD Xtreme™ Hot Start Polymerase with 5% DMSO and 0.3 μM primer using the primers F1 and P7 (24) (**Table S1**). Cycling conditions were as follows: 95°C for 90 sec; 5 cycles of 95°C for 15 sec, 68°C for 30 sec, and 72°C for 30 sec; 13 cycles of 95°C 15 sec, 55°C 30 sec and 72°C 30 sec, followed by a final extension at 72°C for 5 min. Samples were treated with ExoSAP-IT™ PCR product cleanup reagent (Thermo Fisher Scientific) according to manufacturer’s protocol. A second PCR step was performed with the transposon specific primer containing the adapter sequence using KOD Xtreme™ Hot Start Polymerase with 5% DMSO and 0.3 μM primer in a 62.5-μL reaction volume using the primers Tn5-left and P7 (24) (**Table S1**). Cycling conditions were as follows: 95°C for 3 min, 12 cycles of 95°C for 30 sec, 55°C for 30 sec, and 72°C for 30 sec, followed by a final extension at 72°C for 5 min. Product size (~200 bp) was confirmed on a 1% agarose gel. After standard bead cleanup and Illumina library preparation, samples were sequenced at the University of Chicago Genomics Facility using a custom sequencing primer (24) (**Table S1**).

Fitness analysis was performed as previously described (25) using the TRANSIT software (available at https://github.com/mad-lab/transit). We used the permutation test in TRANSIT to quantify differences in sequencing read counts between our PYE and lake water conditions (25). The complete Tn5 dataset (**Tables S4 & S5**), genes with differential fitness (p<0.01; **Tables S5 & S6**), and genes shared between the Tn5 and Tn-Himar datasets (**Table S7**) are listed in Supplemental Information. Raw Tn-seq data are deposited in the NCBI sequence read archive under BioProject accession PRJNA429486; BioSample accession SAMN08348191; SRA accession SRP128742.

## RESULTS

### Growth of Caulobacter in natural freshwater

As a prerequisite to measuring strain fitness, we first sought to demonstrate *Caulobacter* growth in natural freshwater. We collected nearshore water from Lake Michigan, representing a typical oligotrophic freshwater system inhabited by *Caulobacter* spp. (26, 27). With no additional supplementation, filtered (0.1 μm) lake water supported *Caulobacter* growth to a maximal density of approximately 5 × 10^5^ CFU/mL (**Fig. 1A**), from an initial inoculum of 2.5 × 10^4^ CFU/mL. *Caulobacter* doubled 4–5 times at a rate of 0.14 hr^-1^ (doubling time 5 hr). Similar growth rates were observed in unfiltered lake water. Supplementation with 0.1% xylose increased the maximal density by about 10-fold, while addition of 23 mM K2HPO4 had no effect (**Fig. 1A**), implying that carbon, but not phosphorus, limits *Caulobacter* growth in Lake Michigan water. For comparison, we also assayed *Caulobacter* growth in water collected from Lake Superior and found a similar growth yield (**Fig. 1B**). Supplementation with either 23 mM K2HPO4 or 0.1% xylose did not significantly enhance *Caulobacter* growth, but together 0.1% xylose and 23 μM K2HPO4 enhanced growth by more than 10-fold, suggesting that both carbon and phosphorus limit growth in Lake Superior. By comparison, *Caulobacter* reached a density of 3 × 10^9^ CFU/mL in PYE broth or in defined M2X medium (**Fig. 1C**). Notably, cell density was stable for one week in lake water but declined by 2-3 orders of magnitude after 2 days of cultivation in artificial media (**Fig. 1**). This finding is consistent with a report by Poindexter describing *Caulobacter* isolates that tolerated prolonged nutrient scarcity with little loss of viability (11). Based on our results, we chose to perform our genetic analysis in unsupplemented water from Lake Michigan.

**Figure 1.**
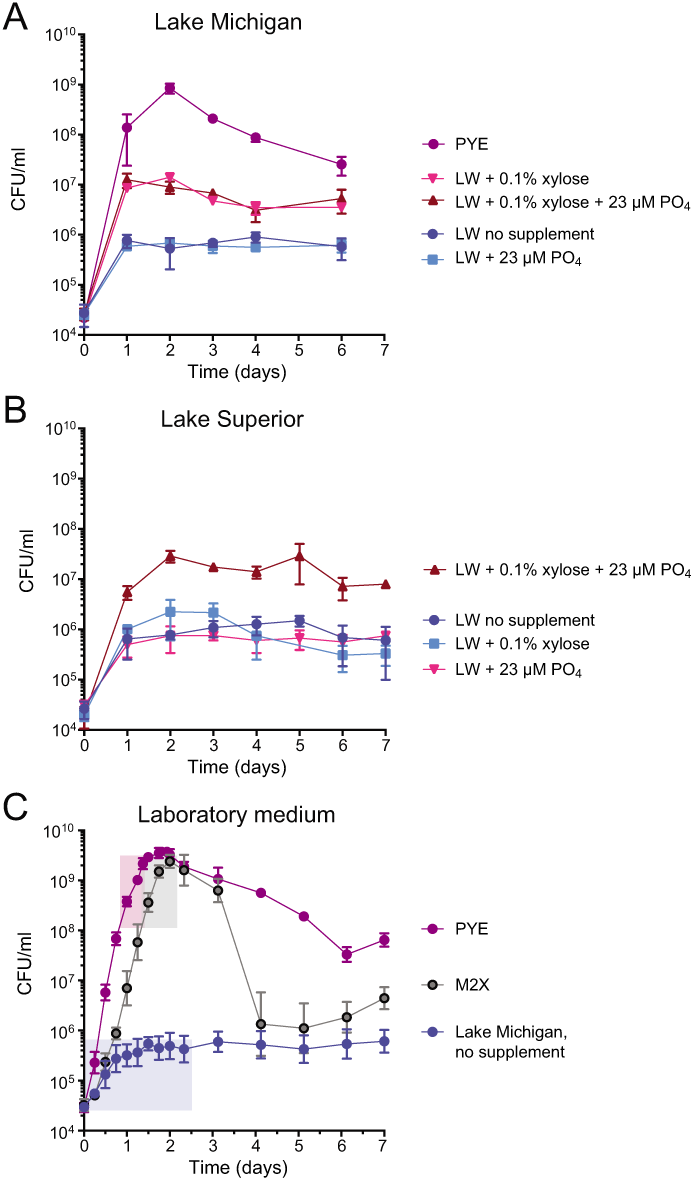
Growth of *Caulobacter* in laboratory medium, and supplemented or unsupplemented water from two Great Lakes. Overnight cultures washed with filtered lake water (LW) were inoculated into 5 mL of filtered water from Lake Michigan (A) or Lake Superior (B). Growth assays in water supplemented with carbon (0.1% w/v xylose) and/or phosphorus (23 μM K2HPO4) as indicated; growth was monitored every 24 hours by enumeration of colony forming units (CFUs) by dilution plating. Lake water growth is compared to growth in a laboratory peptone yeast extract (PYE) medium. Data represent mean ± standard deviation of 5 replicates per condition. (C) Fine scale growth of *Caulobacter* in PYE, M2-xylose defined medium (M2X), and filtered Lake Michigan water. Cells were grown as in A and B and monitored by enumerating CFUs. Data represent mean ± standard deviation of 5 replicates per condition. Boxes represent the approximate region of the growth curve (cell density and incubation time) in which the barcoded Tn-Himar mutant library was cultivated.

### A global Tn-sequencing approach identifies Caulobacter mutants with altered fitness in lake water

We sought to identify genes required for *Caulobacter* growth in natural freshwater, compared to defined M2X or complex PYE medium. To this end, we constructed mutant libraries (**Table 2**) using two different transposons: Tn5, which inserts randomly, and Tn-Himar, which inserts specifically at TA dinucleotides, which occur on average every 82 bp in the *Caulobacter* genome. Each transposon in the pool of Himar transposons contains a unique 20-bp barcode sequence, which is mapped once to a specific insertion site in the genome and thereafter can be quantified by simple amplicon sequencing (9), see **Fig. S1**. Both transposon libraries were constructed by growing cells in PYE; hence insertions in genes essential for growth in PYE are not represented in either library.

**Table 2:** Transposon library statistics

| Library | Unique insertion sites | Total TA sites | Percent sites hit | Average transposons per ORF | Mean reads |
| --- | --- | --- | --- | --- | --- |
| Tn5<sup>a</sup> | 115,788 | N/A | 2.9 | 30 | 90–150 (per Tn) |
| Tn-Himar<sup>b</sup><br>(BarSeq) | 46,395 | 49,437 | 93.8 | 24 | 7612 (per gene) |

<sup>a</sup>Tn5 can insert into any dinucleotide site. We identified 115,788 insertion sites in an initial population of ~300,000 clones.
<sup>b</sup>TnHimar specifically inserts into TA dinucleotides.

We cultivated the *Caulobacter* Tn5 pool in PYE and filtered lake water (0.1 μm). Although Tn5 is capable of insertion at almost every position in the genome, our Tn5 library had lower site saturation than our Tn-Himar library, which limited the statistical power to identify significant fitness effects (25, 28). We calculated mutant fitness and gene essentiality for all genes (**Tables S5 & S6)** and identified 55 genes for which Tn5 disruption significantly diminished or enhanced growth in lake water relative to PYE (adjusted p-value cutoff < 0.01). Given the limited power of the Tn5 dataset, we focused our analyses on the Tn-Himar dataset, but include the Tn5 data in the supplemental material as they provide useful validation of the Tn-Himar data discussed hereforward.

The *Caulobacter* Tn-Himar library contained an estimated 2 × 10^6^ clones, of which 7 × 10^4^ passed the criteria for barcode mapping (9). Considering there are only ~ 5 × 10^4^ TA insertion sites in the *Caulobacter* genome, it is clear that in this population we hit some sites more than once with unique barcodes. We cultivated this library in four conditions: 1) complex PYE medium, 2) defined M2X medium, 3) filtered lake water, and 4) unfiltered lake water (**Fig. S2**). To ensure that we started the experiment with sufficient mutant strain diversity, we inoculated the same total number of cells (2.5 × 10^7^) in each treatment and aimed for 4–5 doublings into the late exponential phase of growth (**Fig. 1 and S2**). For PYE and M2X treatments, cells were grown in 1.5 mL volumes for 10 and 20 h, respectively. For lake water treatments cells were grown in three flasks each containing 2.5 L for 64 hours. By varying culture volume, we ensured an equal number of cell divisions across a similar phase of the growth curve. This approach required cultivation at different cell densities between conditions. After harvest, barcodes were analyzed as described (9), and strain fitness was calculated as the log2 of the ratio of barcode abundance in lake water (or M2X) to the control condition (PYE) (9). Given the 20–30-fold increase in cell number of the mutant pool, a Tn-Himar insertion strain that did not grow at all should have a fitness score around -4 to -5; more extreme (lower) fitness scores indicate strains that did not survive cultivation. The most extreme negative fitness scores in this dataset (i.e. < -4) likely reflect genes that are essential in a particular condition (**Fig. 2A & 2D; Table S2**). The distributions of fitness scores in defined medium and the two lake water conditions are presented in **Figs. 2B & 2C**.

To validate our approach, we examined the fitness consequences of disrupting xylose utilization genes in the M2-xylose (M2X) growth condition. Genes in the *xylXABCD* operon are required for xylose utilization (29, 30). As expected, insertions in these genes generated fitness scores of -3.6 to -6.6 when the pool was cultivated in M2X (**Fig. 3A, Table S2**). Disruption of *xylR*, which functions as a transcriptional repressor of the xylose operon (29), resulted in a positive fitness score in M2X relative to PYE, indicating that derepression of the xylose utilization genes is advantageous when xylose is the sole carbon source. Disruption of the *xylXABCD* genes had little effect on fitness in lake water, which contains a range of carbon sources beyond xylose; disruption of *xylR* resulted in a modest fitness decrease in lake water relative to PYE (**Fig. 3A)**, suggesting a cost to constitutive expression of unused genes.

**Figure 2.**
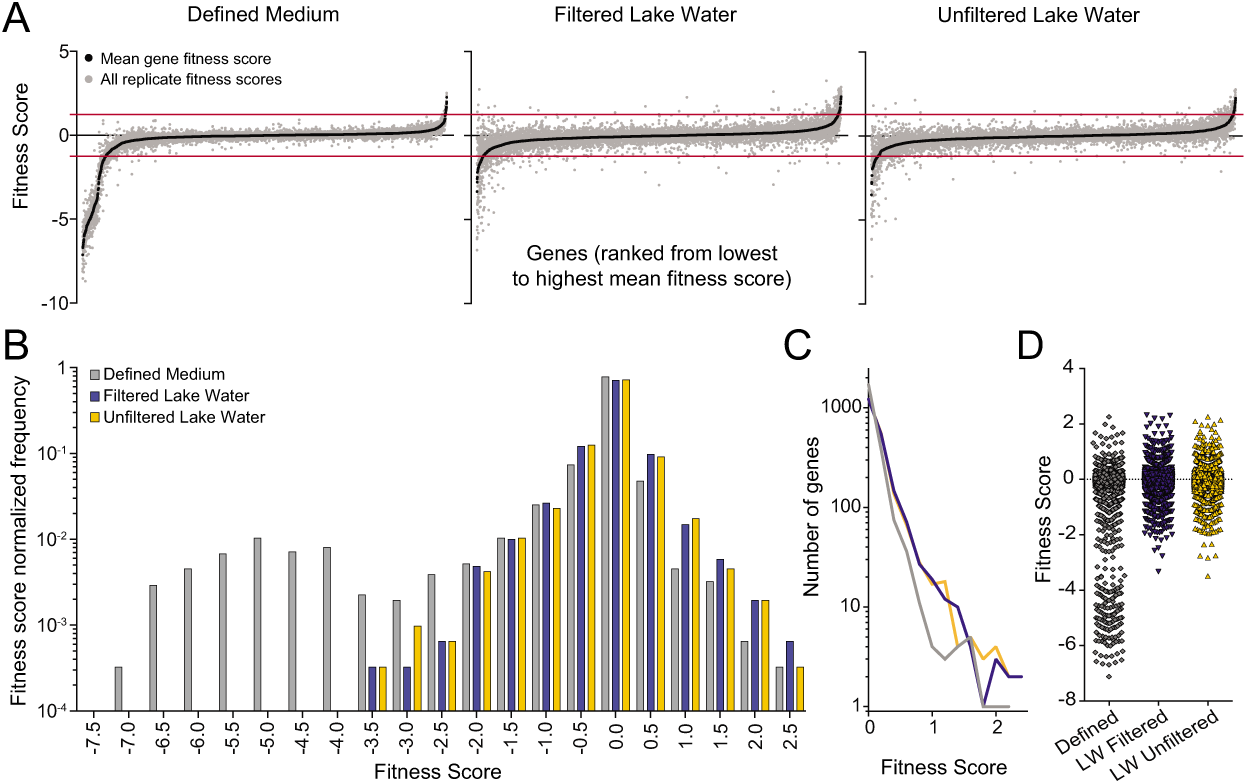
Caulobacter gene fitness score summary after cultivation in defined medium, filtered, or unfiltered Lake Michigan water. (A) Rank ordered mean fitness scores of each scorable *Caulobacter* gene across each of the four replicate experiments for each growth condition is plotted; black = mean fitness score; gray = independent replicate fitness scores. Red lines represent ± 3σ from the mean score of the entire dataset (which is approximately zero; genes with fitness scores less than -2.5 in M2X were excluded from this determination). (B) Distributions of mean gene fitness scores for each condition: defined M2X medium (gray), filtered Lake Michigan water (blue), and unfiltered Lake Michigan water (yellow). (C) Distribution of mean gene fitness values between 0 and +2.5 plotted for each condition; defined M2X medium (gray), filtered Lake Michigan water (blue), and unfiltered Lake Michigan water (yellow). (D) Genes fitness score distribution scores plotted for each of the three cultivation conditions: defined M2X medium (gray), filtered Lake Michigan water (blue), and unfiltered Lake Michigan water (yellow).

**Figure 3.**
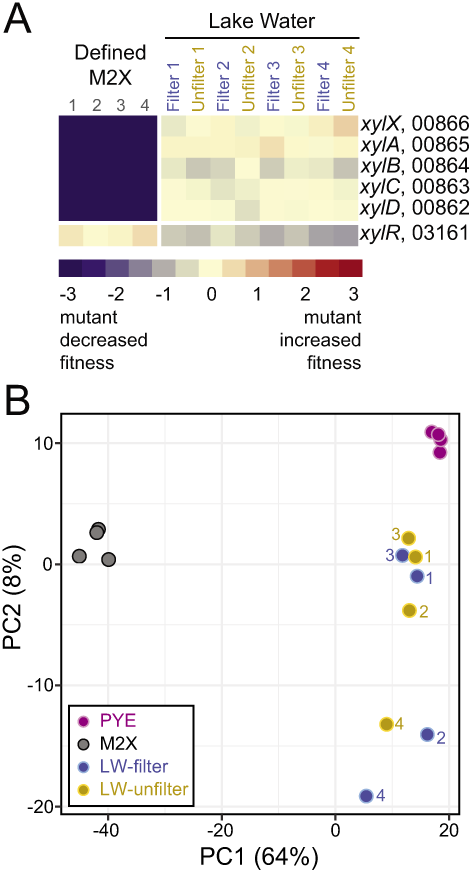
Functional validation of the barcoded Tn-Seq approach; PCA profile of Tn strain fitness in complex medium, minimal defined medium and Lake Michigan water. (A) Heatmap of gene fitness scores for six xylose utilization genes (*xylX, xylA, xylB, xylC, xylD, xylR*) from each replicate experiment of cultivation of the Tn-Himar mutant library in M2X defined medium, filtered lake water, and unfiltered lake water. Fitness score color scale bar is shown below the panel. (B) Principal component (PC) analysis (PCA carried out in ClustVis (49)) of genomic-scale fitness values for the barcoded *Caulobacter* Tn-Himar mutant library cultivated in complex peptone yeast extract (PYE) medium (reference set), defined M2X medium, filtered Lake Michigan water, and unfiltered Lake Michigan water. The plot shows PCA values for individual samples from each cultivation condition. Percent of variance contributed by the first two PCs is noted on the axes. PCA plot is based on all fitness score values for all genes in the Tn-Himar datasets (see Table S2).

### Increased variability of fitness scores in lake water

Compared to defined M2X medium, lake water is more heterogeneous over time and space. Our four lake water experiments used water collected on four days over a 2-week period and showed greater variability in strain fitness scores than our four independent M2X replicates (**Figs. 2A, 3B & 4**). In addition, we fit a linear model to test the effects of two factors, sampling day and filtration condition, on strain fitness scores, and found a number of genes that differed significantly across days, including genes related to cell surface carbohydrate biosynthesis and environmental sensing and gene regulation (**Table S8**). This variability likely reflects day-to-day differences in temperature, mixing, and biotic factors such as phage dynamics, though we cannot completely rule out technical day-to-day variations in sample processing. Future work with additional temporal replicates could discriminate genes whose functions are consistently important from genes that are exploited under transient conditions in the lake.

Surprisingly, filtration (0.1μm) had little effect on the global fitness profile of *Caulobacter* (**Fig. 4**), and our linear model approach did not identify any genes with differential fitness between filtered and unfiltered lake water. This result implies that particulates and other microorganisms present in the lake water did not affect strain growth, suggesting that *Caulobacter* is not in strong competition with other microbes for “common goods” in this system on the time scale of our experiment.

**Figure 4.**
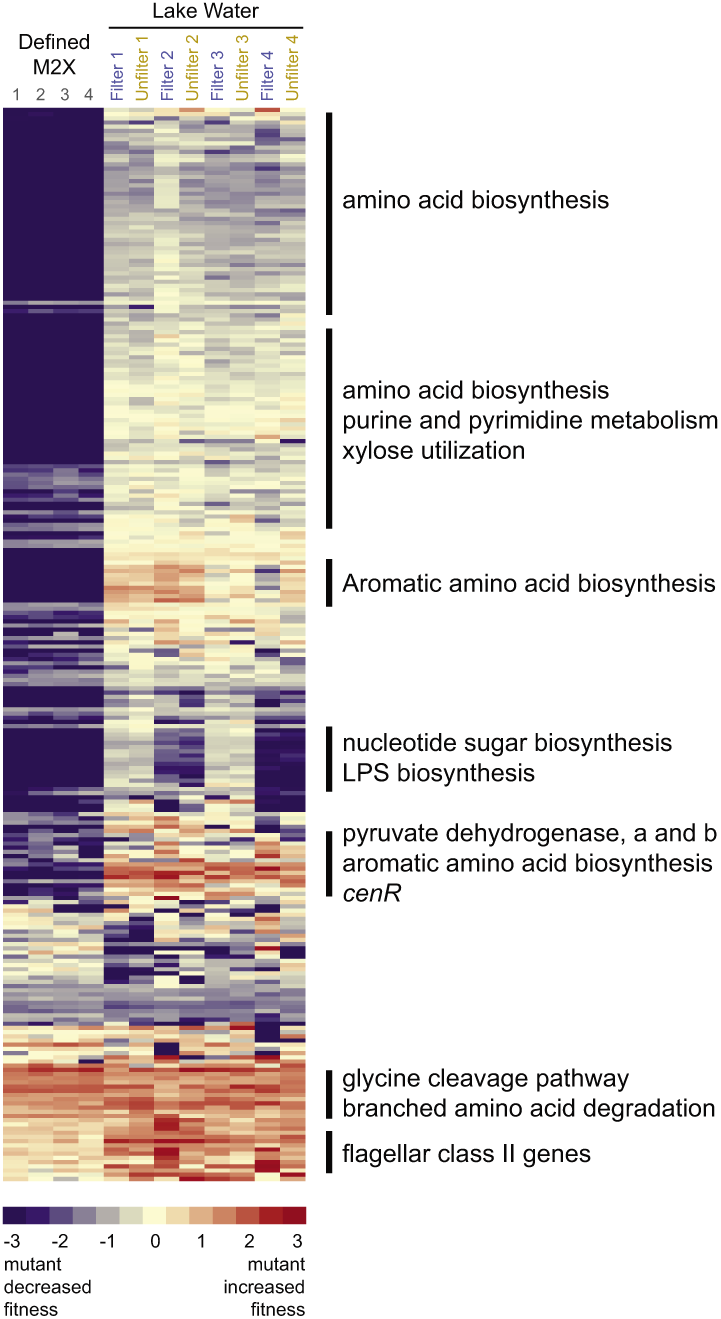
Functional summary of mutant strains with diminished or enhanced fitness in minimal defined medium and Lake Michigan water. Heatmap of fitness scores for genes with mean fitness scores higher than 1.2 or lower than -1.2 (approximates ± 3σ fitness score cutoff) in at least one cultivation condition. The sigma cutoff was based on the major fitness score distribution centered on zero (see Figure 2B). Genes are hierarchically clustered using Cluster 3.0 (average linkage) and visualized using TreeView, and fitness scores for each replicate experiment are color coded on scale bar shown below. General functions of genes within particular regions of this 256 gene cluster are noted; this entire figure is expanded and split into three fully annotated clusters in Fig. S5, with gene names included. The full cluster of genes with mean fitness scores higher than +1.2 or lower than -1.2 in the lake water conditions (excluding the M2X data) is presented in Fig. S6.

### Fitness defects are more extreme in defined medium than in lake water

Transposon disruption of genes required for amino acid biosynthesis, nucleotide biosynthesis, lipopolysaccharide biosynthesis, and nucleotide sugar biosynthesis resulted in extreme (fitness score < -4) growth defects in M2X (**Fig. 3, Tables S2–S3**); these fitness scores provide evidence that strains harboring disruptions of these genes did not grow at all in M2X and thus likely comprise a strain/gene set that is essential in this defined condition. This result is not surprising, considering that growth in M2X medium requires de novo biosynthesis of diverse monomers and intermediates, many of which are supplied exogenously in the reference PYE condition. In many cases, strains with severe fitness defects in M2X also had reduced growth in lake water, but the fitness costs were less severe (**Tables S2 & S9-S10**). We controlled the number of doublings (approximately 4–5) across all conditions, so the more pronounced fitness costs in defined medium compared to lake water cannot be explained by differences in the number of doublings. Instead, these results imply that lake water is more similar to the reference condition PYE than M2X is to PYE, in terms of the metabolic demands it imposes on cells. This inference is supported by principal component analysis across all growth conditions (**Fig. 3B**). Indeed, we expect that natural freshwater supplies diverse metabolites and growth substrates that may render some genes dispensable, whereas defined media provides fewer exogenous resources.

### Pathways conferring differential fitness in natural freshwater and artificial media

To further explore the selective pressures faced by *Caulobacter* across these conditions, we focused on genes whose disruption induced large fitness effects, namely fitness scores less than -1.2 and greater than +1.2 (this approximates a ±3s cutoff). Based on this criterion, we identified 83 and 82 genes in the filtered and unfiltered lake water conditions, respectively, and 213 genes in the defined M2X medium (**Table S9**). Genes with significant fitness values across all three conditions based on the t-statistic of Wetmore and colleagues (9) are outlined in **Table S10**. Broad functional patterns in our Tn-Himar dataset were assessed using clusters of orthologous group (COG) annotations (31) (**Fig. 5**). A full comparison of genes for which Tn-Himar disruption results in a specific advantage or disadvantage in M2X defined medium, but not in filtered or unfiltered Lake Michigan water (relative to complex PYE medium), and vice versa, are presented in **Tables S11-S12**. Genes that were not hit by Tn-Himar, and thus not included in any of our analyses are included in **Table S13**. Many of these genes have been previously defined as essential (18). A clustered heatmap that contains genes with fitness scores less than -1.2 and greater than +1.2 from either the filtered or unfiltered lake water conditions is presented in Fig. S6.

**Figure 5.**
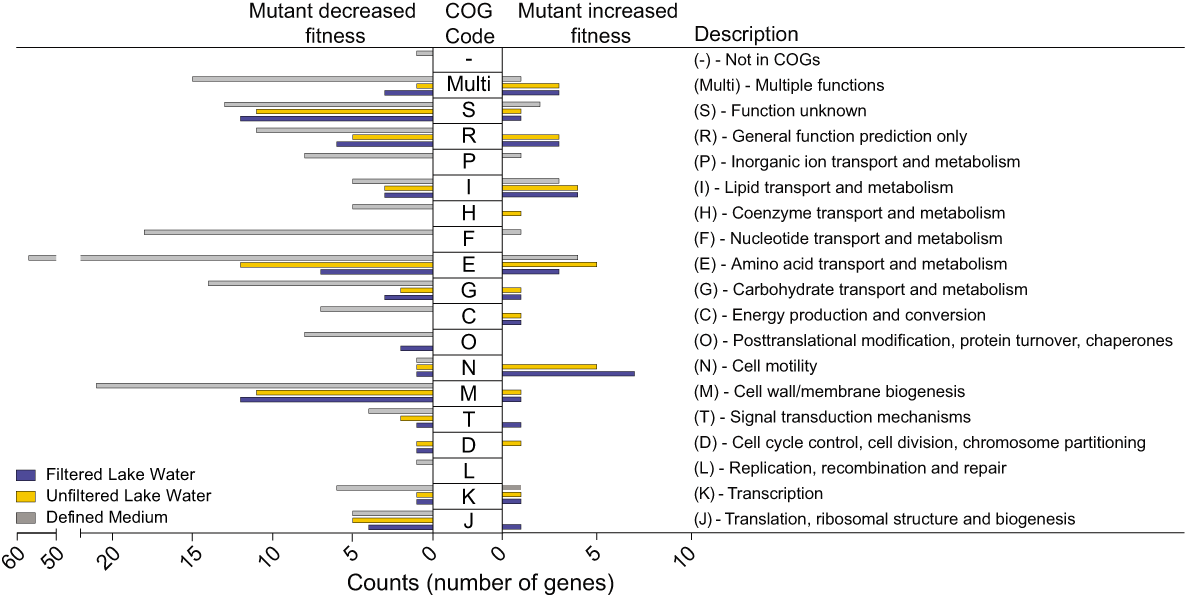
COG analysis of Tn-Himar gene fitness data. The analysis includes genes with fitness scores of absolute value greater than 1.2 (which approximates a ± 3σ fitness score cutoff) in each condition. Each gene was assigned a cluster of orthologous group (COG) functional category, obtained through the NCBI COG site (31). The number of genes in each COG category is plotted; genes with negative fitness values (left) and genes with positive fitness values (right).

Not surprisingly, the most negative fitness scores were observed for genes in amino acid and nucleotide biosynthesis (discussed above), and in genes required for transport of metabolic substrates into the cell (**Tables S2 and S9**); we observed similar defects in our Tn5 dataset (**Tables S4-S6**). In addition, disruption of genes encoding catabolic enzymes in the glycine cleavage pathway and in branched amino acid degradation led to an apparent enhancement of fitness in both M2X medium and in lake water relative to PYE, in both the Tn-Himar and Tn5 experiments (**Fig. 4; Tables S2 & S4-S7**). This result likely reflects the higher cost of deleting these catabolic genes in the reference PYE condition compared to M2X or lake water, and is consistent with transcriptional data showing that select amino acid degradation pathways — including glycine cleavage, histidine, branched chain, and phenylalanine degradation — are upregulated in PYE compared with M2X (32).

Surprisingly, we found enhanced fitness for strains with disruptions in motility genes in lake water relative to PYE (**Fig. 4 & 5**). We more carefully examined the fitness scores of genes involved in synthesis and assembly of the flagellum (**Fig. 6**). The flagellum is assembled in a regulated hierarchy of stages, which is well described in *Caulobacter* (33-35). Class II genes encode the inner components of the flagellum, including the export apparatus, and regulatory proteins that activate expression of class III and IV genes. Class III genes encode the basal body and hook structures. Completion of class III structures activates translation of class IV genes, which encode the subunits of the flagellar filament. Thus, defects in each class prevent expression of subsequent classes. Within each class of flagellar genes, we observed consistent fitness patterns, demonstrating the power of this method to capture even modest effects of gene disruption. Disruption of class II flagellar genes conferred an advantage that was significantly greater in lake water than in M2X compared to PYE (**Fig. 6B & S2**). Disruption of class III genes followed similar trends, but with smaller magnitude effects. *Caulobacter* encodes six redundant class IV flagellin genes (36), three of which are represented in our Tn-Himar pool and whose disruption, not surprisingly, had no effect on fitness. Disruption of the motor stator gene *motA* or *motB*, which results in a fully assembled but paralyzed flagellum (37, 38), did not affect fitness under our cultivation conditions. Together, these results suggest that the fitness advantage of flagellar gene disruption is not derived from energy saved in powering the flagellum, but rather in energy or resources saved in synthesizing and assembling the flagellum. In the lake water cultivations, we observed appreciable day-to-day variation in the fitness of each class of flagellar gene mutants (**Fig. S2**), which was particularly pronounced for class III genes. Patterns in this day-to-day variability were consistent across members of each class, suggesting that this variability is driven by environmental factors rather than technical factors.

**Figure 6.**
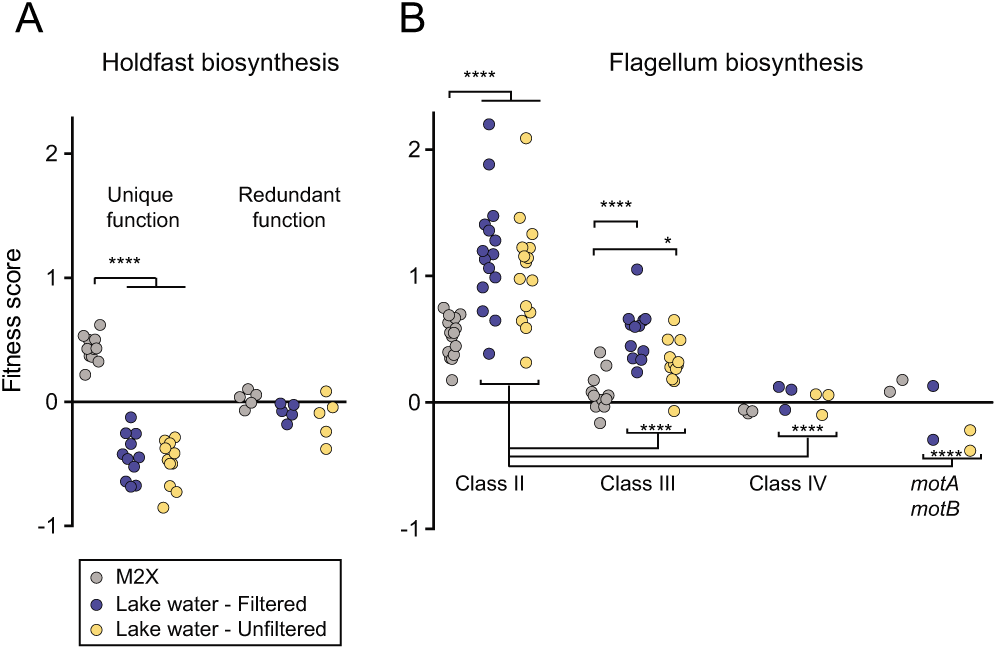
Genes with functions in flagellum and holdfast biosynthesis influence fitness in Lake Michigan water. Mean fitness scores of genes involved in (A) holdfast biosynthesis (17 genes) and (B) flagellum biosynthesis (38 genes) are plotted for each cultivation condition (four independent growth replicates): defined M2X medium (gray); filtered lake water (blue); and unfiltered lake water (yellow). Holdfast genes with unique function (i.e. single gene deletions have a holdfast defect) and redundant function (i.e. single mutants have no holdfast defect) are shown separately. Class II, Class III, and Class IV genes in the flagellar hierarchy and *motA*/*motB* stator complex genes are also shown separately. Clusters of holdfast and flagellum fitness score data, with individually annotated genes, are shown in Fig. S4. One-way ANOVA was applied to assess differences in fitness scores between marked groups; Tukey’s post test (**** p < 0.0001; * p < 0.05).

Fitness was also affected by the ability to synthesize the polar extracellular adhesin known as the holdfast (**Fig. 6A & S2**). We systematically analyzed genes involved in synthesis, secretion, and attachment of the holdfast. Most holdfast genes yield partial to complete defects in holdfast development when disrupted (39); we categorized these genes as ’unique functions’ genes. However, two sets of holdfast biosynthesis genes have redundant functions: two Wzy-family polymerase genes function in holdfast development and three paralogs of the HfsE glycosyltransferase have genetically redundant activities in holdfast synthesis (39). Disruption of genes in these redundant sets had no effect on fitness. Disruption of genes in the unique function group resulted in a modest but consistent fitness advantage in M2X and a fitness disadvantage in both filtered and unfiltered lake water, relative to PYE (**Fig. 6A & S2A**). For the group of all unique genes, the fitness consequence for loss of holdfast was significantly different between growth conditions (p < 0.0001) (**Fig. 6A**).

### Genes of unknown function contribute to fitness in natural freshwater

We hypothesized that many genes of unknown function play important roles in natural environmental contexts but not in typical laboratory media. Of all genes showing large fitness effects (± 3σ), hypothetical genes or genes of unknown function accounted for 16% (13/83) in filtered lake water, 15% (12/82) in unfiltered lake water, and 7% in defined medium (15/213) (**Table 3**). Across these three conditions, five hypothetical genes were shared. *CCNA_03860* was the only hypothetical gene for which disruption provided a fitness benefit across all three conditions relative to PYE. CCNA_03860 contains a conserved domain belonging to the YkuD superfamily, which has been shown to have L,D-transpeptidase catalytic activity, providing an alternate pathway for peptidoglycan cross-linking (40, 41). Disruption of *CCNA_01724, CCNA_03864, CCNA_03909*, and *CCNA_00375* resulted in reduced fitness across all three conditions relative to PYE. Hence using natural growth conditions may be critical for understanding the functions of many uncharacterized bacterial genes.

**Table 3:** Average fitness scores for hypothetical genes and genes of unknown function with fitness scores less than -1.2 and greater than +1.2 (bold and shaded) in at least one condition. LW, Lake Michigan water.

| Locus number |  | Defined | Filtered LW | Unfiltered LW |
| --- | --- | --- | --- | --- |
| CCNA_00375 | No conserved domains | **-1.21** | **-2.57** | **-2.83** |
| CCNA_01724 | COG4649, TPR_21 pfam09976 | **-2.58** | **-1.25** | **-1.28** |
| CCNA_03860 | COG3786, YkuD superfamily | **1.33** | **1.53** | **1.41** |
| CCNA_03864 | DUF3576, pfam12100 | **-1.71** | **-1.32** | **-1.57** |
| CCNA_03909 | No conserved domains | **-1.20** | **-1.76** | **-1.84** |
| CCNA_00927 | No conserved domains | **-2.56** | **-2.11** | -0.67 |
| CCNA_02875 | No conserved domains | **-1.54** | **-2.19** | -0.39 |
| CCNA_00895 | No conserved domains | -0.19 | **-2.14** | **-1.44** |
| CCNA_00913 | No conserved domains | -0.28 | **-2.48** | **-1.21** |
| CCNA_00519 | No conserved domains | **-1.66** | -0.37 | -0.61 |
| CCNA_01176 | DUF2849, pfam11011 | **-4.98** | -0.81 | -0.84 |
| CCNA_01178 | DUF934, pfam06073, COG3749 | **-4.36** | -0.60 | -0.64 |
| CCNA_01676 | TamB, COG2911, pfam04357 | **-6.40** | -0.52 | -0.74 |
| CCNA_01219 | No conserved domains | **1.68** | -0.70 | 0.78 |
| CCNA_02669 | Uncharacterized membrane protein, DUF3422, pfam11902, COG4949 | **-4.41** | 0.38 | 0.20 |
| CCNA_03273 | COG4944, DUF1109, pfam06532 NrsF, anti-sigF | **-1.63** | -0.39 | -0.27 |
| CCNA_03692 | COG1975, XdhC/CoxF family | **-4.06** | -0.98 | -0.65 |
| CCNA_02796 | No conserved domains | -0.45 | **-1.20** | -0.52 |
| CCNA_03420 | No conserved domains | 0.40 | **-1.46** | 0.42 |
| CCNA_03883 | No conserved domains | 0.05 | **-1.25** | 0.99 |
| CCNA_03984 | No conserved domains | 0.01 | **-1.22** | -0.25 |
| CCNA_01286 | No conserved domains | 0.25 | -0.25 | **-1.27** |
| CCNA_02053 | No conserved domains | -0.56 | -0.74 | **-1.34** |
| CCNA_02160 | No conserved domains | 0.26 | -1.02 | **-1.38** |
| CCNA_03015 | No conserved domains | -0.15 | 0.57 | **-1.46** |
| CCNA_03945 | No conserved domains | -0.43 | 0.24 | **-1.90** |

LW, Lake Michigan water.

## DISCUSSION

### Tn-seq fitness scores provide a window into cell-environment interactions

Bacterial genomes carry relatively little noncoding DNA. Genes that confer no fitness benefit tend to decay over time (42) implying that genes that are maintained are beneficial at least under some circumstances. Yet traditional microbial cultivation approaches often fail to yield discernable mutant phenotypes for many genes. One approach to overcome this challenge is to interrogate gene function in more relevant ecosystem contexts, embracing physicochemical complexity. The genome-scale fitness analysis of *Caulobacter* transposon mutants reported in this study provides new understanding of genes that affect growth in a *bona fide* freshwater environment. Disruption of genes involved in biosynthesis of non-aromatic amino acids, lipopolysaccharides, and nucleotide sugars results in large fitness defects in natural freshwater compared to complex laboratory medium (PYE). Moreover, fitness effects were variable across temporal lake water replicates; this variability likely reflects physicochemical and biological variability in the lake and suggests an important role for transient response genes in fluctuating environments.

### The fitness costs and benefits of motility and attachment in freshwater

The energetic cost of flagellar biosynthesis and motility is well established (43, 44). Our data indicate that transposon disruption of genes required for the synthesis of the single polar flagellum of *Caulobacter* enhanced fitness in lake water relative to PYE medium (**Fig. 6B**). This is consistent with a *Salmonella* Tn-Seq study that revealed a fitness advantage in strains with disrupted flagellar genes (45). Notably, we found that fitness effects were not uniform across all flagellar genes: disruption of class II genes, which has the greatest impact on flagellar gene expression, also led to greater effects on fitness, compared to class III and IV genes. The fitness enhancement in lake water is not due to the energy savings from motor rotation, as strains with insertions in the *motA* and *motB* stator genes, which assemble a full but non-rotary flagellum (38), showed no fitness difference (**Fig. 6B).** We conclude that the relative fitness advantage of flagellar gene disruption is related to the cost of biosynthesis of flagellar proteins. It seems certain that over longer cultivation timescales, and in more spatially complex environments, the *Caulobacter* flagellum provides a fitness advantage, as flagellar genes are maintained in natural freshwater environments.

Our data reveal that disruption of genes required for holdfast biosynthesis is disadvantageous when strains are cultivated in lake water relative to PYE. This fitness cost was evident in both filtered (particle-free) and unfiltered lake water relative to PYE (**Fig. 6A**), suggesting that the effect is not due to adhesion to particles in the medium. Instead, it is possible that the holdfast confers a growth advantage by enabling adherence to the flask surface, where polymeric nutrients concentrate to form conditioning films (46, 47). In defined M2X medium, disruption of holdfast biosynthesis genes confers a fitness advantage (**Fig. 6A**). In this medium, all the components are salts or simple sugars, which do not efficiently condition naïve surfaces (46, 47). In this case, surface attachment is apparently not advantageous, and holdfast biosynthesis comes at a cost.

### Genetic evidence suggests a complex medium is a better freshwater analog than a defined mineral medium

Fitness defects of *Caulobacter* mutants were often more severe in a defined mineral xylose medium (M2X) than in lake water, relative to PYE. Moreover, the overall fitness profile of *Caulobacter* mutants cultivated in lake water more closely resembles that in PYE than in M2X, suggesting that dilute complex medium is a better proxy for natural freshwater. *Caulobacter* belongs to a group of dimorphic prosthecate (i.e. stalked) alphaproteobacteria that are often specialized for oligotrophic, dilute environments (10, 11). Indeed, the inhibition of growth and stalk development due to excess nutrients was the first physiological property of *Caulobacter* spp. to be described (48). Complex and defined media of varying compositions have been outlined for cultivation of *Caulobacter* and related genera, but it is notable that dilute peptone (less than 0.2% w/v) generally supports growth of all dimorphic prosthecate bacteria (11). This observation supports the notion that the natural nutrient environment of this class of bacteria is best captured by cultivation in a dilute complex medium that contains amino acids and other trace complex biomolecular components. Our data also demonstrate that an M2-based medium exerts highly specific metabolic constraints and is likely not an ecologically or physiologically relevant growth condition.

### An approach to study gene function in ecosystem context

The explosion of bacterial genome sequence information has far outpaced our ability to characterize gene function using traditional approaches, leading to the accumulation of thousands of ’unknown’ protein families. Many of these families are conserved throughout the bacterial domain, which is evidence that they confer a selective benefit in particular conditions. This leads to the following question: under what circumstances do these conserved families provide a fitness advantage? At the onset of this study, we hypothesized that many of these unknown protein families would prove to be important in the natural ecological context of a bacterium. Among the genes whose disruption leads to the greatest fitness effects (± 3σ) in filtered lake water relative to PYE, approximately 15% are hypothetical or conserved genes of unknown function (**Tables 3, S8–S9**). The approach we describe here indicates that these genes of unknown function play an important role in *Caulobacter* physiology in a natural freshwater environment. Going forward, one can take advantage of lake-specific growth phenotypes to begin to define the functions of these genes in an ecologically relevant context.

## ACKNOWLEDGEMENTS

The authors have declared no competing interests. This work was supported by UChicago BIG grant to S.C. and M.C, and NIGMS grant R01GM087353 to S.C. K.L.H was supported by an NIH Ruth Kirschstein Postdoctoral Fellowship (F32 GM122242) and a Chicago Biomedical Consortium Postdoctoral Core Grant (FP064244-01-PR). L.R.R was supported by the NIH Molecular and Cellular Biology Training Grant (T32 GM007183). P.D.C. is supported an NSF CAREER award (1552647); he began the Tn5 library construction in the laboratory of Dr. Yves V. Brun at Indiana University. We thank the members of the Crosson laboratory for helpful discussions, Tom Ioerger (Texas A&M) for assistance with TRANSIT, Adam Deutschbauer (University of California-Berkeley) for the *E. coli* APA752 strain, and David Hershey for construction of the *Caulobacter crescentus* CB15 Himar transposon library. We also thank Pieter Faber and Abhilasha Cheruku from the University of Chicago Genomics Facility for technical advice and helpful discussions.

